# PcoB is a defense outer membrane protein that facilitates cellular uptake of copper

**DOI:** 10.1101/2021.11.26.470079

**Authors:** Ping Li, Niloofar Nayeri, Kamil Gorecki, Eva Ramos Becares, Kaituo Wang, Dhani Ram Mahato, Magnus Andersson, Sameera Abeyrathna, Karin Lindkvist-Petersson, Gabriele Meloni, Julie Winkel Missel, Pontus Gourdon

**Affiliations:** Lund University Department of Experimental Medical Science, Sölvegatan 19, SE-221 84, Lund, Sweden; University of Copenhagen, Department of Biomedical Sciences, Nørre Allé 14, DK-2200, Copenhagen N, Denmark; Department of Chemistry, Umeå University, Linneaus Väg 10, 901 87 Umeå, Sweden; Department of Chemistry and Biochemistry, The University of Texas at Dallas, 800 W Campbell Rd., Richardson, TX 75080, USA

**Keywords:** Outer Membrane Protein Structure, Gut microbiota, PcoB

## Abstract

Copper (Cu) is one of the most abundant trace metals in all organisms, involved in a plethora of cellular processes. Yet elevated concentrations of the element are harmful, and interestingly prokaryotes are more sensitive for environmental Cu stress than humans. Various transport systems are present to maintain intracellular Cu homeostasis, including the prokaryotic plasmid-encoded multiprotein *pco* operon, which is generally assigned as a defense mechanism against elevated Cu concentrations. Here we structurally and functionally characterize the outer membrane component of the Pco system, PcoB, recovering a 2.2 Å structure, revealing a classical β-barrel architecture. Unexpectedly, we identify a large opening on the extracellular side, linked to a considerably electronegative funnel that becomes narrower towards the periplasm, defining an ion conducting pathway as also supported by metal binding quantification via ICP-MS and MD simulations. However, the structure is partially obstructed towards the periplasmic side, and yet flux is permitted in the presence of a Cu gradient as shown by functional characterization *in vitro*. Complementary *in vivo* experiments demonstrated that isolated PcoB confers increased sensitivity towards Cu. Aggregated, our findings indicate that PcoB serves to permit Cu import. Thus, it is possible the Pco system physiologically accumulates Cu in the periplasm as a part of an unorthodox defense mechanism against metal stress. These results point to a previously unrecognized principle of maintaining Cu homeostasis and may as such also assist in the understanding and in efforts towards combatting bacterial infections of Pco-harboring pathogens.

## Introduction

Copper (Cu) is a transition metal essential for virtually all organisms, for example serving as a co-factor for a number of enzymes involved in redox reactions. However, elevated Cu levels is associated with mismetallation and damage to proteins and cells, and catalyze toxic reactive oxygen and nitrogen species production via redox cycling (2). Strikingly, mammals are frequently more tolerant to increased Cu levels in the surroundings than prokaryotic counterparts (3). Organisms have developed mechanisms for tight regulation of the Cu levels (4). The significance of maintained Cu homeostasis is underscored by the many different protein networks linked to this process. In *Escherichia coli,* the cytoplasm is maintained devoid of free Cu via its export mediated by the Cue/Cop system, regulated by the transcription factor CueR (5) (**Figure 1**, **yellow** (6, 7)). CopA, an inner membrane P-type ATPase, extrudes Cu^+^ ions to the periplasm (8), where it is oxidized to less toxic Cu^2+^ by CueO, a multicopper oxidase (9). When this response is overwhelmed or under anaerobic conditions, when the CueO oxidase is inactive, the Cus-mediated Cu export assembly is activated (**Figure 1, magenta** (10)). Cus connects the inner and outer membranes, spanning the entire periplasm (10) through three proteins CusCBA, and is energized by the tripartite resistance-nodulation-cell division (RND) CusA component, collectively providing capacity to export Cu^+^ from the cytoplasm directly out of the cell (11). Additionally, CusF, a periplasmic Cu-sequestering protein, delivers the metal directly to CusB for efflux (12). The expression of the Cus constituents is regulated by CusRS (13, 14). Considering the Cus system limitations, complementary Cu homeostasis proteins exist, most notably the plasmid-born Cu resistance Pco system (**Figure 1, cyan**). This operon was first detected in *E. coli* from the gut flora of pigs fed at high Cu diet (15); Cu in combination with antibiotics have been used as growth promotor in pig diets for at least 45 years (16).

**Figure 1.**
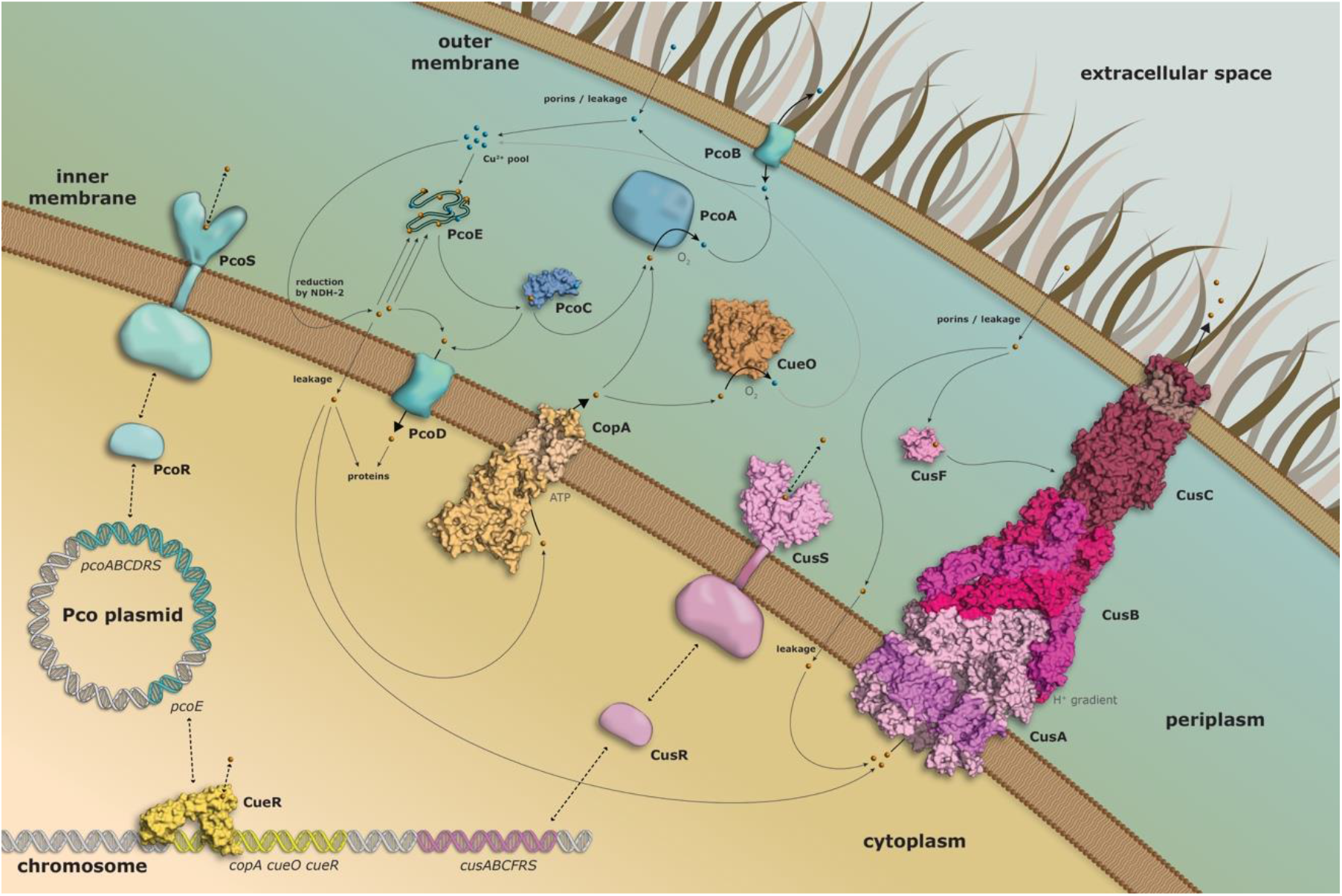
Overview of copper homeostasis proteins in gram negative bacteria. The Cue/Cop system (in yellow) is responsible for copper detoxification through removal from the cytoplasm via CopA and periplasmic oxidation via CueO as regulated by CueR. At higher copper concentrations or anaerobic conditions, the Cus assembly (purple) provides export from the cytoplasm via CusA, and from the periplasm via CusF, immediately to the extracellular environment as allowed by CusB and CusC and with the expression regulated by CusR and CusS. The plasmid born Pco cluster (cyan) likely has a complementary role, harboring an inner and outer membrane component, PcoD and PcoB, respectively, and the periplasmic oxidase PcoA as well as the copper-binding PcoC and PcoE proteins, as regulated by PcoR and PcoS.

The Pco proteins have been shown to enable bacteria to survive at higher Cu concentrations, although *E. coli* strains lacking the genes accumulates less Cu in the periplasm and exhibit higher Cu efflux (15, 17). Underscoring the significance of the Pco assembly, homologous proteins are frequently present on chromosomes or plasmids of other bacteria, where they also have been linked to increased Cu tolerance (15, 18). Nonetheless, there is growing evidence congruent with the Pco proteins also being involved in Cu uptake (17, 19), seemingly in conflict with the observed role for Cu defense. Thus, even the physiological role of the Pco proteins for Cu homeostasis in bacteria remains enigmatic.

The pco gene cluster in *E. coli* encompasses seven genes, pcoABCDRSE (20). The PcoRS is a two-component regulatory system, analogous to CusRS, sensing the periplasmic Cu concentrations (20). PcoE resides in the periplasm and binds Cu, predicted to serve as a ‘molecular sponge’, thereby decreasing the free Cu concentration in the compartment between the two cell membranes (21). PcoD represents an inner membrane protein, and the function is likely tightly linked to periplasmic PcoC as they often exist as a fusion protein. For example, the *Bacillus subtilis* single protein YcnJ shares high sequence homology to the two PcoCD components. Deletion of YcnJ is associated with impaired growth in low Cu media suggesting a putative role in Cu acquisition (17), while expression of PcoABD leads to Cu hypersensitivity in the absence of PcoC (22).

Similarly to PcoCD, PcoAB have been proposed to work together as the primary actors in pco-dependent Cu resistance (23). While PcoA is a periplasmic multicopper oxidase, distantly related to CueO, PcoB resides in the outer membrane and has an elusive function (20). PcoB has been suggested to prevent Cu uptake from the cellular outside (24), however, since CopA appears necessary for Pco-dependent Cu resistance, PcoB is generally believed to be a Cu-specific transport protein, acting in concert with PcoA (24). Homologues of PcoAB are regularly encoded in close proximity in Gram-negative bacteria, and whereas PcoA is sometimes found alone, PcoB is always accompanied by PcoA, suggesting the interaction between the two and that PcoB requires PcoA for the Cu-transport function (25). For instance, expression of PcoB alone in the absence of PcoA in a ΔpcoAB *Caulobacter crescentus* strain did not rescue the Cu-sensitive phenotype (26). However, PcoC was also shown to be needed for full resistance of the Pco system, and to interact with PcoA, possibly serving as a periplasmic Cu-chaperone (27). Collectively, the molecular details of the function and the regulation of the Pco system remains elusive.

In this work, we set out to elucidate the physiological role and functional properties of PcoB. We determine the 3D structure and characterize the protein function *in vitro* and *in vivo*. Our findings shed fundamentally new light on the role of the Pco system in Cu homeostasis.

## Results and Discussion

### Pco confers cell survival at elevated Cu levels but isolated PcoB sensitizes to Cu stress

To further dissect the physiological role of the Pco system, we first compared *E. coli* cells transfected with a vector harboring the operon or a similar control vector lacking the Pco components. Using electron microscopy, the strains were studied at low and high Cu levels, respectively. While in a Cu-low environment both systems emerged as healthy intact cells (**Figure 2A upper panels**), the strain without the Pco system appeared impeded under Cu stress (**Figure 2A left bottom panel**). In contrast, cell viability was maintained at high Cu concentrations for the cells with the *pco* operon (**Figure 2A right bottom panel**), yet displaying a somewhat different morphology compared to the cells proliferated in Cu-depleted conditions. Specifically, the Pco harboring cells preserve the overall shape and display separation of the cell interior from the plasma membrane, while many cells without the Pco system have adopted an elongated character, with certain cells even appearing disrupted. Collectively, these findings suggest the Pco proteins serve a role for cell survival at elevated Cu concentrations, congruent with the bulk of the literature on the system. Nonetheless, the more rarely detected, somewhat contracting, observation of Pco-mediated import is puzzling (28), and highlights the enigmatic role of PcoB, representing the first line of defense from the outside. Is it facilitating import or export of Cu?

**Figure 2.**
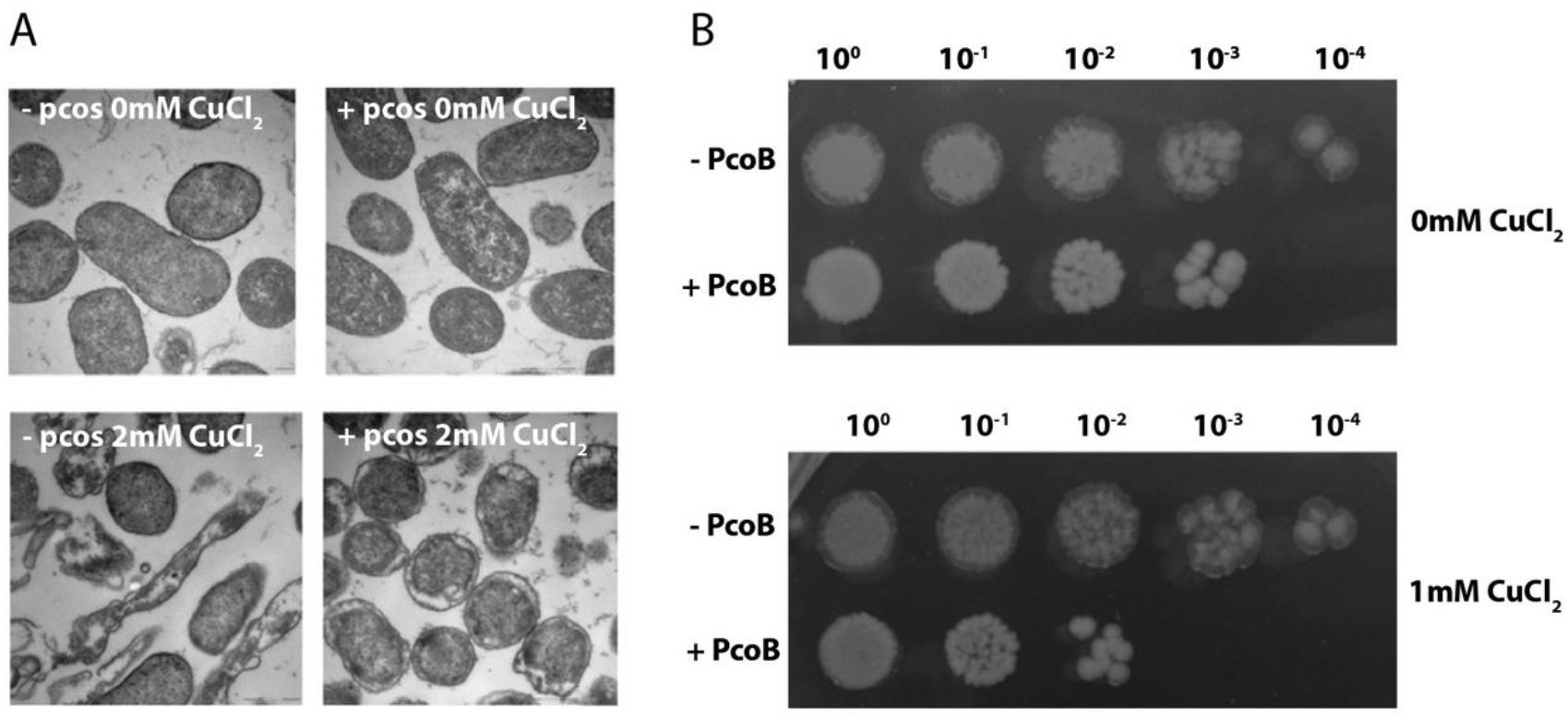
*In vivo* experiments support a protective role of the Pco system while isolated PcoB increases copper sensitivity. (A) Introduction of the Pco gene cluster (right panels, labelled with +) rescues *E. coli* viability at elevated (2 mM) copper concentrations (lower panels), as compared to cells lacking the Pco system (left panels) (B). Comparison of the growth of *E. coli* cells with or without isolated PcoB at low (no supplementation) and high (1 mM) copper concentrations, respectively, suggests PcoB alone increases the copper susceptivity of cells

Next, to shed further light on the specific function of PcoB, we investigated the protein outside of the context of the other Pco proteins. Surprisingly, *E. coli* cells containing PcoB showed increased sensitivity towards Cu compared to cells without the protein in a growth assay (**Figure 2B**). Thus, PcoB appears to serve as an importer, alternatively other components of the Pco system are necessary for the system to operate as an exporter, as CusA is required for CusC.

### The molecular structure of PcoB

To elucidate underlying mechanism behind the observed contradicting physiological responses, we sought out to obtain high-resolution structural information of PcoB. The protein was overproduced, extracted from *E. coli* outer membranes and purified to near homogeneity, eluting at a size-exclusion chromatography (SEC) retention volume that indicated monomeric particle distribution. However, efforts to crystallize full-length PcoB, PcoBFL, were fruitless. In contrast, N-terminally truncated PcoB, PcoB_Δ32-82_ (**Figure 3 - figure supplement 1**), a form that does not alter the Cu-binding properties (**Figure 3A**), successfully yielded crystals in the presence of the detergent C_8_E_4_, which diffracted to 2 Å. Nevertheless, a reliable molecular replacement solution was not identified. Instead, the structure was determined using single-wavelength anomalous diffraction (SAD) based on SeMet-data (**Table 1**) and refined to R/Rfree=0.22/0.26. Overall, the generated electron density maps are of high-quality, permitting assignment of individual sidechains (**Figure 3B**), with the exception for the N-terminus and a single loop (residues Gly82-Ala89 and Asp230-Arg238), which appears disordered (**Figure 3 - figure supplement 2**).

**Figure 3.**
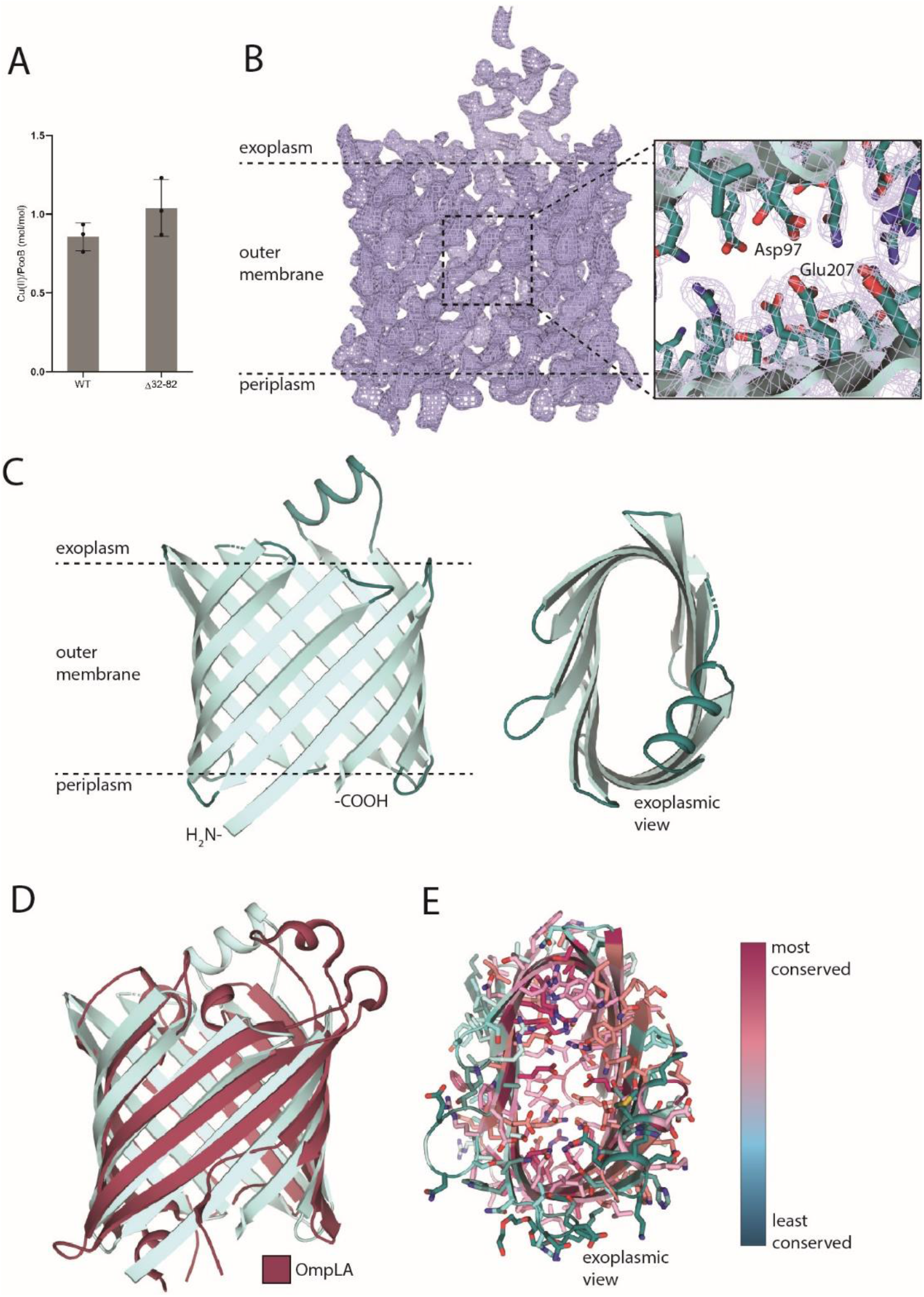
The structure of PcoB. (A) Copper binding stoichiometry as determined using ICP-MS measurements of full-length (wild-type) and N-terminally truncated PcoB, respectively. A single Cu^2+^ ion binds per PcoB molecule. Data denotes 3 independent experiments measurements (N = 3) and error bars represent SD. (B) Final 2Fo-Fc electron density (blue mesh, σ = 1.0) of PcoB derived from the 2.2 Å native data. The close view demonstrates the general quality of the map and the Asp97-Glu207 interaction in the pore region. (C) The overall architecture of PcoB (in cyan) consisting of 12 beta-sheets that span the outer membrane. The PcoB barrel (cyan) is flattened through interactions between residue of opposite sides of the inside of the barrel, see also panel D. (D) Overlay with structurally reminiscent protein OmpLA (dark red, PDB-ID: 1QD6). (E) Analysis of the conservation of PcoB as determined using ConSurf (31). The analysis shows a spectrum ranging from low (cyan) to high (magenta) conservation, as also depicted by the color bar. Internal residues are generally more conserved than membrane facing residues.

**Table 1.** Crystallographic table of PcoBΔ32-82 for both native and SeMet crystals.

|  | <u>NATIVE</u> | <u>SE-MET</u> |
| --- | --- | --- |
| **<u>DATA</u>** |  |  |
| COLLECTION WAVELENGTH | 1 | 0.979587 |
| RESOLUTION RANGE | 43.53 – 2.0 (2.072 – 2.0) | 46.25 - 2.62 (2.74 - 2.62) |
| SPACE GROUP | C 2 2 2<sub>1</sub> | P 4<sub>1</sub> 2<sub>1</sub> 2 |
| UNIT CELL | 65.49 Å 75.54 Å 91.51 Å<br>90° 90° 90° | 64.76 Å 64.76 Å 198.20 Å<br>90° 90° 90° |
| TOTAL REFLECTIONS | 103621 (10567) | 343385 (41634) |
| UNIQUE REFLECTIONS | 15638 (1528) | 13396 (1554) |
| MULTIPLICITY | 6.6 (6.9) | 25.6 (26.8) |
| COMPLETENESS (%) | 99.72 (99.74) | 99.7 (98.2) |
| MEAN I/SIGMA(I) | 16.51 (2.11) | 19.0 (0.8) |
| R-MERGE | 0.06975 (0.8315) | 0.103 (5.697) |
| CC1/2 | 0.999 (0.8) | 1.0 (0.676) |
| CC* | 1 (0.934) | 1.0 (1.0) |
| ANOMALOUS COMPLETENESS |  | 99.7(98.1) |
| ANOMALOUS MULTIPLICITY |  | 14.3(14.4) |
| **<u>REFINEMENT</u>** |  |  |
| REFLECTIONS USED IN REFINEMENT | 15631 (1525) |  |
| R-WORK | 0.2205 |  |
| R-FREE | 0.2533 |  |
| NUMBER OF NON-HYDROGEN ATOMS | 1686 |  |
| MACROMOLECULES | 1599 |  |
| LIGANDS | 40 |  |
| SOLVENT | 47 |  |
| RMS (BONDS) | 0.008 |  |
| RMS (ANGLES) | 1.08 |  |
| AVERAGE B-FACTOR | 48.02 |  |
| MACROMOLECULES | 47.97 |  |
| LIGANDS | 51.31 |  |
| SOLVENT | 46.87 |  |

The structure discloses a classical β-barrel (**Figure 3C**), formed by 12 antiparallel strands that span the outer membrane *in vivo*. The termini are located at the same end of the barrel, in agreement with periplasmic localization, as established by numerous studies on proteins with a barrel topology (29). The strand linking loops are generally short, except for a loop with a helix insertion, stretching into the extracellular space (**Figure 3C**). Notably, the overall fold is reminiscent to that of the outer membrane phospholipase A1 (OmpLA), despite low sequence similarity and unique molecular functions (**Figure 3D**). Indeed, OmpLA contains extensive loops blocking the orifice, and a hydrophobic interior (30), characteristics that separate the protein types. However, the architecture of the PcoB barrel is flattened as a consequence of inter-barrel contacts between charged amino acids, thus restricting the width (**Figure 3C** and **E** and **Figure 4D** and **E**), whereas these interactions of OmpLA are caused by hydrophobicity. The majority of the sheets consist of alternating hydrophobic and hydrophilic residues, facing the surrounding environment and pointing into the channel, respectively, as typical for outer membrane proteins. Equivalently, sequence (Clustal Ω) and surface (ConSurf) (31) conservation analyses suggest residues with sidechains located inside the β-barrel to be relatively conserved, while membrane exposed amino acids are not (**Figure 3E**, **Figure 3 - figure supplement 3**). Thus, it is conceivable that PcoB operates as a monomer, as also supported by the single molecule observed in the asymmetric unit of the crystal packing.

**Figure 4.**
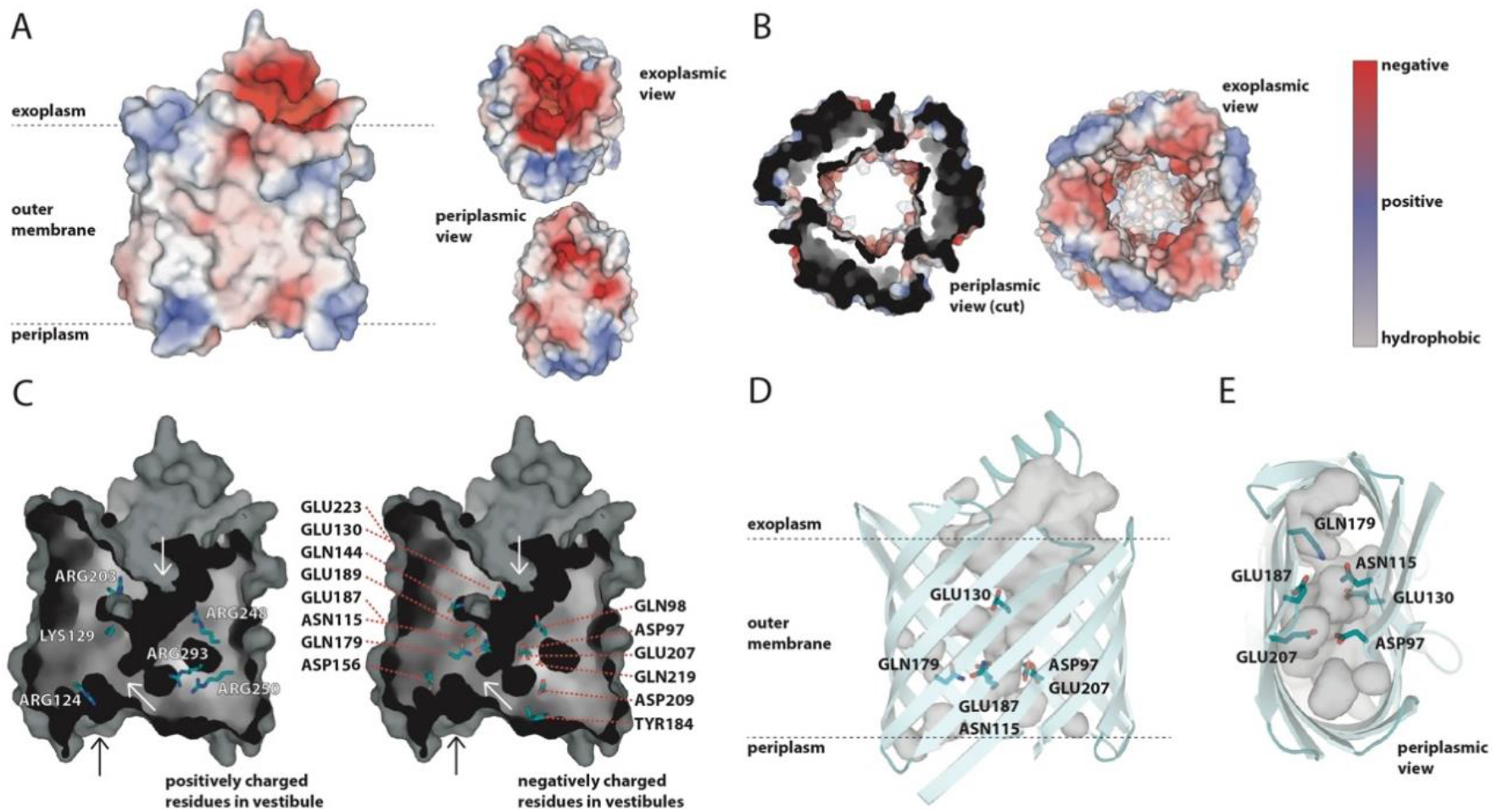
The pore. (A) Surface charge representation of PcoB as observed perpendicular to the membrane, complemented by views from the outside of the cell and from the periplasm, respectively. Electrostatics are represented as positive (blue), negative (red) and neutral (grey) charges. (B) Electrostatic representation of CusC (PDB-ID: 3PIK) using the same color representation as in panel A. The periplasmic view is cut at the outer membrane interface, removing the soluble domain. (C) Surface representation of the pore and internal cavities combined with residues of positive and negative charge, respectively. The top white arrow points to the funnel, the middle white arrow points to the internal constriction site. The black arrow indicates the vestibule observed in panel A bottom view. (D) Side-view perpendicular to the membrane. Important residues for the here proposed copper conducting pathway are shown. (E) Periplasmic view (same orientation as the periplasmic view in panel A), showing the same features as panel D.

### PcoB harbors a highly electronegative pore compatible with Cu uptake

Strikingly, a 12×18 Å wide cavity is present on the extracellular side (**Figure 4A**, top right), as established by condensed loops between strands in the region leaving the aperture uncovered. From this region, a funnel-like pore (**Figure 4C-E, Figure 4 - figure supplement 1A**) is penetrating almost the entire barrel, becoming narrower towards the periplasm, with a remarkably electronegative funnel surface throughout (**Figure 4A**, bottom right). The outline of the pore is supported by the presence of numerous and continuous crystal waters (the maximum observed oxygen-to-oxygen distance is 4.4 Å) (**Figure 4 - figure supplement 1A**). Along the pore, multiple negatively charged-paired residues are present, in particular: Glu130-Glu223 towards the extracellular side, Asp97-Glu207 and Asn115-Gln179 towards the periplasm, which are both placed perpendicular to Glu187 (**Figure 4C-E**). The Glu130-Glu223 and Asp97-Glu207 pairs are both highly conserved among PcoB proteins, and form interacting carboxylic acid-carboxylate hydrogen bonds that assist in flattening the barrel, and at the same time lines the pore. This negatively charged interior reminds of the outer membrane protein component, CusC, of the CusABC system (**Figure 4B**) (32). The pore properties and the similarity to CusC are congruent with PcoB metal conductance as Cu2^+^ rather than Cu^+^. This is because, based on the HSAB Pearson theory (33), Cu^+^ typically rely on soft ligand coordination (Met and Cys) for ion transfer, while the Cu2^+^ is favored by funneling and potential coordination by harder ligands (such as Asp, Glu or His). The presence of these potential transient ligands in the pore hints at inward Cu2^+^ flux (from the outside), considering the highly electronegative surface on the extracellular side, presumably attracting divalent ions from the outside of cells. However, the structure was obtained in Cu-free conditions, and consequently there are no indications of the metal in the pore. Analogously, soaking and co-crystallization experiments to identify Cu presence in the pore were fruitless. In contrast, inductively coupled plasma mass spectrometry (ICP-MS) data on purified protein suggest Cu2^+^ is bound in the pore when metal has been supplemented to the sample as PcoBFL and PcoB_Δ27-81_ each display one bound Cu2^+^ per molecule (**Figure 3A**). Corresponding analysis using ICP-MS of the of the Asp97Lys mutant indicates a significantly impaired Cu^2+^ binding to PcoB (approximately 70 % lower than PcoBFL) in agreement with the Asp97-Glu207 being directly involved in high affinity Cu binding at the end of the funnel or that the anticipated salt-bridge in the mutant prevents Cu passage, closing the pore (**Figure 5**, **Figure 4 - figure supplement 1**).

**Figure 5.**
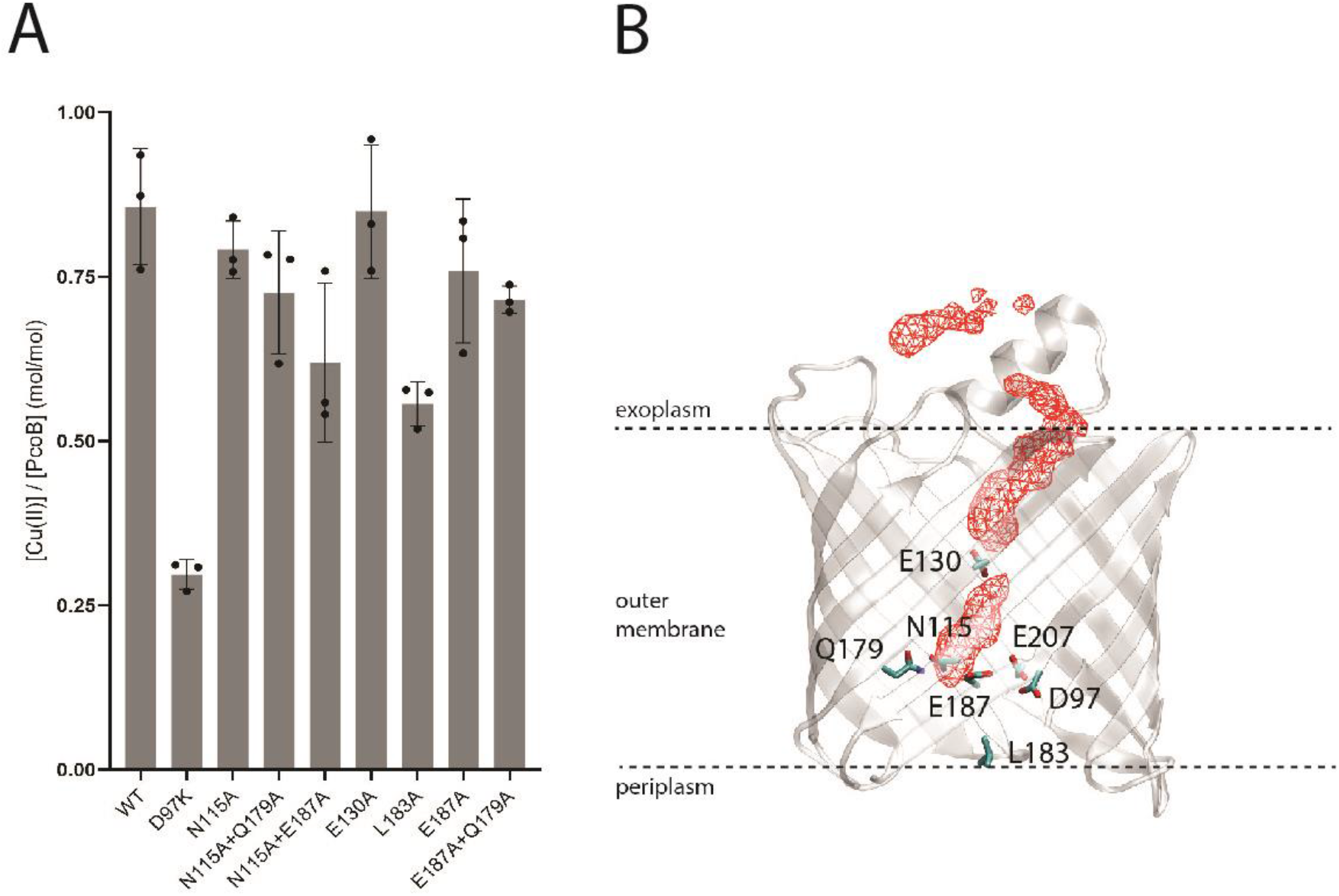
Restricted passage and copper binding. (A) Cu^2+^ binding stoichiometry of PcoB mutants as determined using ICP-MS. Data denotes 3 independent measurements (N = 3) and error bars represent SD. The data is congruent with a single ion binding site located at the Asp97-Glu207 constriction. (B) Isodensity surface (red) at 70 % occupancy representing Cu^2+^ positions in the MD simulation to assess ion passage across PcoB.

In this light, we set out to investigate if PcoB indeed facilitates Cu^2+^ flux across cellular membranes. Utilizing a protocol developed for reconstitution and flux measurements of other porins (34), PcoBFL and the Asp97Lys mutant were successfully reconstituted, and protein-free liposomes employed as a as control (**Figure 6**). As evident from the averaged traces from three reconstitutions, wild-type clearly conducts Cu^2+^ (**Figure 6A**), while the mutation appears less active than the wild-type (**Figure 6B**), suggesting the central Asp97 represents a key feature for permitting Cu binding and flux.

**Figure 6.**
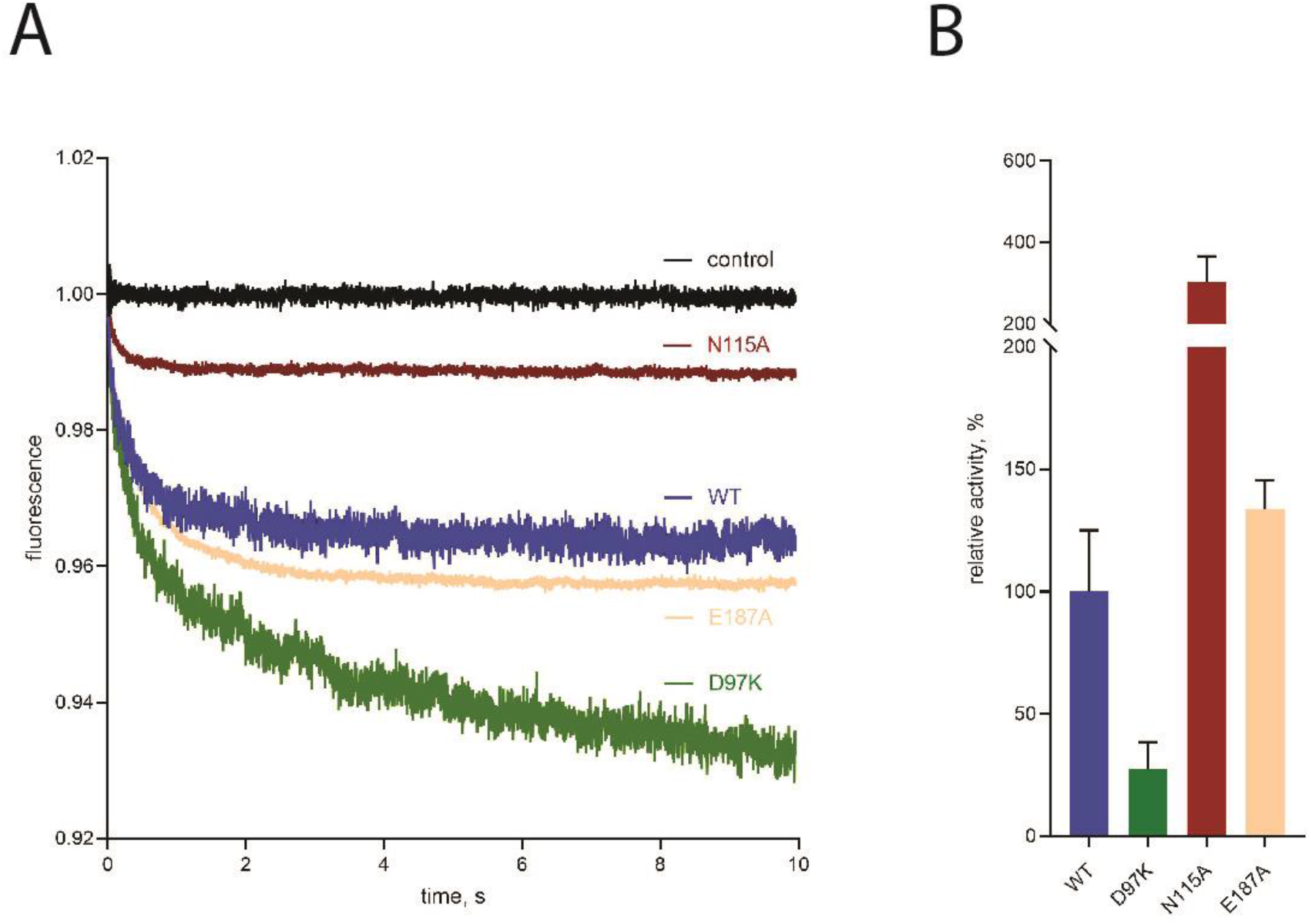
Functional characterization based on a proteoliposome assay. (A) Wild-type PcoB (blue) demonstrates flux of Cu^2+^, clearly deviating from the control measurements (black) performed using empty liposomes. Data are also shown for the three assessed mutant forms. Traces originate from 5 runs based on triplicate reconstitutions. (B) Bar diagrams of the relative activity of the investigated PcoB mutants with wild-type set as 100 %. The activity has been compensated for the amount of protein incorporated into the liposomes (**Figure 6 - figure supplement 1, Figure 6 - table supplement 1**). The shown forms were all well-behaved throughout the isolation and characterization process. Data are means +− SD of the three separate reconstitutions (N = 3) for each construct.

### The periplasmic constriction

In contrast to the extracellular side, the waterline of the pore becomes partially obstructed at the intersection between Asp97, Glu187, Asp209 and Glu207, with Asp209 and the adjacent Tyr184, being in proximity to the periplasm and yet capping the pathway together with Leu183 (**Figure 4C, Figure 4 - figure supplement 1**). Through analysis of the water molecules in the region detected in the structure, two possible Cu2^+^ paths are revealed towards the periplasm (**Figure 4C** and **Figure 3A**), as also confirmed by surface analysis of the internal cavities. However, one of these is lined by several positive residues, most notably Arg250 and Arg293 in the immediate vicinity of Asp97-Glu207, disfavoring involvement in Cu passage (**Figure 4C**). In contrast, the second path is rich in negatively charged residues (**Figure 4C**), and links directly to the electronegative periplasmic vestibule (**Figure 4A**) through a network consisting of Asn115, Gln179 and Glu187 (**Figure 4C-E**) Notably, these three possible gate residues are well conserved (Asn115 is frequently replaced by other long-side chain residues) and, in addition, analysis of evolutionary coupled pairs using the EVcouplings server (https://evcouplings.org/) suggest Gln179 to be linked to Asn115, in agreement with a shared functional purpose (**Figure 4 - figure supplement 1**) (35). Alanine replacements of these residues had little to no effect on the binding stoichiometry as evaluated by ICP-MS experiments (**Figure 5**). This may suggest these amino acids are rather important for the overall Cu conductance when the pore is fully open, without establishment of discrete sites that would affect binding stoichiometry.

Considering we were unable to obtain a complementary structure in the presence of Cu, to further investigate how ions may bridge the restriction region, we instead generated alanine substitutions *in silico* of relevant residues to assess the effect on the local environment. Notably, mutations of Leu183, Tyr184 and Asp209 on the Arg250-Arg293 pathway do not result in a significant pore widening **(Figure 4 - figure supplement 1C**), despite the presence of crystal waters along this pathway. On the other hand, replacements of the residues of the complementary vestibule, with Glu187, Asn115 and Gln179, resulted in a significant reduction of the constriction independent of mutation (**Figure 4 - figure supplement 1B**). Notably, mutation of Gln179 alone caused a substantial widening, hinting at the Asn115-Gln179-Glu187 vestibule pathway playing an important role for the Cu^2+^ passage (**Figure 4 - figure supplement 1B**). Next, unbiased molecular dynamics simulations of the recovered PcoB structure were exploited to shed further light on the ion passage. Congruent with our structural analysis, we observed that Cu^2+^ is not capable of passing through this constriction towards the periplasm (**Figure 5B**). Such behavior has however also been found for certain well-established ion channels such as ClC-1 (36), where a narrow constriction region is bridged by ion-transferring sidechains, representing a possible role of Asn115-Gln179-Glu187 in PcoB.

We then reverted to our liposome assay to investigate wild-type and mutant PcoB forms *in vitro* (**Figure 6**). Indeed, alanine substitution of Asn115 and Glu187 with the intention to widen the pore elevated the Cu^2+^ flux. The Asn115 form showed a dramatic three-fold increase of the transfer rate, while Glu187Ala was augmented by 50%, congruent with a role in gating of the ion transfer. Two double mutations of the constriction region, Asn115Ala/Gln179Ala and Glu187Ala/Gln179Ala, however yielded an additional band on SDS-PAGE following liposome reconstitution. We interpreted the response as a sign of instability and refrained from further analysis of these mutant forms (**Figure 6 - figure supplement 1**, **Figure 6 - table supplement 1**, **Figure 6 - source data 1**). It is possible that the shift of Gln179 is responsible for the apparent degradation. Another mutant form in this region, Asp156Lys, demonstrated aggregation already in the size-exclusion purification (**Figure 6 - figure supplement 2**), perhaps due to the change of the local charge.

Analogously, alanine replacement of Leu183, a surface exposed residue that is capping the Arg250-Arg293 pathway, also suffered from instability as detected by SDS-PAGE. Tyr184Ala and Asp209Lys, the latter in immediate connection to Arg250, demonstrated a similar behavior as Asp156Lys, despite Tyr184 being in direct contact with the surrounding environment (data not shown). Thus, it is likely that Leu183, Tyr184 and Asp209 play a role in maintaining the local architecture rather than participating in ion flux (**Figure 6 - figure supplement 1, Figure 6 - table supplement 1**, **Figure 6 - source data 1**). The functional characterization was also complemented by ICP-MS measurements of several of the mutant forms that were possible to recover in detergent form, however the metal binding stoichiometry remain largely unaffected for all forms (**Figure 5**). Thus, it is likely bridging between the extracellularly exposed funnel and the periplasm does not include ion bindings sites per se, rather a direct transfer via amino acid ligands. Aggregated, these findings point towards an expected sensitivity for mutations of the periplasmic-facing region of PcoB. Furthermore, the data suggest ion transfer is achieved via the single Cu^2+^-binding site in the entire protein, located at the very end of the electronegative funnel at Asp97-Glu207 (mutation of which obstructs conductance). Next Cu^2+^ passage occurs through Asn115-Gln179-Glu187 (with pathway enlarging single alanine substitutions elevating the flux), and finally, via the electronegative vestibule that is in direct contact with the periplasm, without presence of distinct check points equivalent to Asp97-Glu187.

### The *pco* operon may act as a defense mechanism in high Cu environments

The composition of the *pco* operon readily suggests its role in maintenance of periplasmic Cu levels: three of the core proteins are periplasmic proteins, responsible for detoxification (PcoA) and Cu binding (PcoC and E) (21, 22) (**Figure 1**). Moreover, the two-component system PcoRS, senses periplasmic Cu levels and regulates the expression of the other proteins, except PcoE that is regulated by CusRS (20). Additionally, the *pco* plasmid has no effect in cells that are unable to export cytoplasmic Cu: deletion of the Cu^+^ exporting P-type ATPase CopA abolishes the protective effect of *pco* (8), an observation that is in agreement with the notion that pco operon is a part of the periplasmic Cu defense mechanism.

The nature of the protection provided by the *pco* operon has remained unclear, yet the functions of three of the proteins are well-established: PcoE being a Cu “sponge”, capable of rapid binding a large number of Cu ions (21), and PcoC being a Cu chaperon that binds both Cu^+^ and Cu^2+^, and delivers Cu^+^ to PcoA, a multicopper oxidase that catalyzes the oxidation of Cu^+^ to Cu^2+^, rendering Cu less toxic (22). In contrast, the specific roles of the membrane proteins PcoB and PcoD have been more elusive. PcoB, a classical outer membrane protein as shown in this work, was previously suggested to provide an efflux pathway for periplasmic Cu, presumably as the last step of a molecular defense system against high environmental stress. However, such efflux requires energy input, as the exported ion moves against a gradient: the extracellular environment contains more Cu than the periplasm when cells are exposed to Cu stress.

Cu export to the extracellular side is achieved in *E. coli* by the CusFCBA system (**Figure 1 in magenta**), the architecture of which is however diametrically different than that of the *pco* system (**Figure 1 in cyan**). CusA is an inner membrane protein that exploits the energy stored in the proton gradient to pump out Cu^+^ ions via a large tunnel consisting of CusB and CusC proteins, together spanning the entire periplasm and forming an exit pathway through the outer membrane (12). The direct coupling between the inner membrane, the periplasm, and the outer membrane allows for the energy-requiring Cu^+^ export reaction to occur. The *pco* system does not contain proteins sufficiently large to form a direct contact to the inner membrane. It is possible that the Cu chaperone PcoC delivers Cu^2+^ to PcoB, which then becomes permeable upon PcoC binding, yet such scenario still does not handle the issues with providing energy for the export against a gradient. These arguments all support the notion of PcoB serving as an importer, although such a function may be counter-intuitive for a defense system.

Upon environmental stress, the outer membrane becomes the first barrier in the defense. Non-specific porins are downregulated and a number of selective outer membrane proteins are rather inserted in the membrane, allowing for selective influx of required molecules while keeping the stressors outside the cell (37). It is possible to imagine a setup where Cu is allowed to leak into the periplasm to acceptable levels (i.e. enough to metallize the selected proteins, but not more). This would require a periplasmic control system, ready to react to elevated concentrations of Cu. A more sophisticated strategy of orchestrating the expression of a Cu-specific porin and downstream Cu chaperones would allow for more control and quicker response.

Our structural, functional and *in vivo* data corroborate the notion that PcoB play a role in such an unorthodox defense system. The structure is highly electronegative and harbors a considerable funnel freely accessible from the outside, clearly in agreement with attraction of charged matter in the surroundings (**Figure 4**). Nevertheless, PcoB is partially obstructed as detected by our structural analysis, suggesting a gradient may be necessary to allow passage. Indeed, Cu flux is permitted when a gradient is applied using the employed proteoliposome assay (**Figure 6**). Analogously, the complete PcoB system increases viability at elevated concentrations of Cu, although the morphology of the cells is altered, perhaps due to the increased levels of metal in the periplasmic space (**Figure 2**). In contrast, PcoB alone does not provide a similar molecular defense and instead increases the metal sensitivity, as the other components of the Pco assembly do not complex the imported Cu.

Taken together, our findings improve our understanding how bacteria handle excess environmental Cu, likely importing Cu^2+^ through PcoB to the periplasm which may serve a dual purpose of providing essential metal at low Cu conditions and/or sequester environmental metal under Cu-stress along generating a pool of periplasmic Cu that can be utilized for cellular import (by PcoB to the periplasm and perhaps also to the cytoplasm via PcoD) during Cu starvation. The results also open up an attractive possibility to increase the Cu sensitivity in PcoB containing pathogens through blockage of the electronegative funnel from the outside of the cells, as a means to combat infections.

## Materials and Methods

### *In vivo* copper tolerance assays

For the agar plate study, bacterial BL21(DE3) cells were transformed with plasmid pET-22b-PcoB and grown on LB-agar plates supplied with 50 μg/mL ampicillin for 16 h at 37 °C (these conditions were used for all copper tolerance growths unless otherwise is stated). Cells harboring empty vector pET-22b were used as control. A single colony was inoculated in 5 mL LB culture for approximately 8 h until the optical density at 600 nm (OD_600nm_) reached 0.8. The cells were spin down, washed and resuspended in fresh LB media to OD_600nm_=0.1, and then diluted in 10-fold increments using LB media. 5 μL drops were spotted onto the LB-agar plates containing defined amounts of CuCl_2_ (0 and 1 mM), and with the media pH-adjusted to 7.0 (following supplementation of CuCl_2_). The plates were incubated at 37 °C for 16h to compare the growth of the colonies.

For the electron microscopy study, the transformed cells were first cultured in 5 mL LB to OD_600nm_=0.6, and then the cells were cultured for 16 h with starting OD_600nm_=0.05 in 5 mL containing the desired CuCl_2_ concentration (0 and 2 mM), with the pH of the media adjusted to 7.0 (following supplementation of CuCl2). The cultured cells were pelleted using 6000 xg for 10 min and then washed three times using PBS buffer to remove dead cells. Pellets of bacteria were fixed with 2 % v/v glutaraldehyde in 0.05 M sodium phosphate buffer (pH=7.2). The pellets were embedded in Agarose, rinsed three times in 0.15 M sodium phosphate buffer (pH=7.2), and subsequently postfixed in 1 % w/v OsO_4_ with 0.05 M K_3_Fe(CN)_6_ in 0.12 M sodium phosphate buffer (pH=7.2) for 2 h. The specimens were dehydrated in graded series of ethanol, transferred to propylene oxide and embedded in Epon according to standard procedures. Sections, approximately 60 nm thick, were cut with a Ultracut 7 (Leica, Wienna, Austria) and collected on copper grids with Formvar supporting membranes, stained with uranyl acetate and lead citrate, and subsequently examined with a Philips CM 100 Transmission EM (Philips, Eindhoven, The Netherlands), operated at an accelerating voltage of 80 kV. Digital images were recorded with an OSIS Veleta digital slow scan 2k x 2k CCD camera and the ITEM software package.

### Expression and purification of PcoB

The gene coding for *E. coli* PcoB including its signal peptide (UniProt Accession No.Q47453) was codon optimized and synthesized by Genscript. A 6xHis tag followed by a TEV cleavage site was introduced between S26 and V27 by overlap PCR to facilitate protein purification. The N-terminus was truncated using PCR, thereby removing 55 residues (27-81) to generate PcoB_Δ27-81_ Next, the full-length PcoB and PcoB_Δ27-81_ were cloned into the pET-22b expression vector using *Nde*I and *Xho*I cleavage sites. The expression plasmids were transformed into the C43 (DE3) *E. coli* strain. Single colonies were incubated at 30 °C for 16 h in 5 mL LB medium supplemented with 100 mg/mL ampicillin. The preculture was transferred into 1 L LB cultures supplied with 100 mg/mL ampicillin and cultivated at 30°C with shaking at 180 rpm. Protein expression was induced at 25 °C for 16 h with final concentration of 0.5 mM Isopropyl β-D-1-thiogalactopyranoside (IPTG) when the OD_600_=0.6.

To solve the crystallographic phase problem via selenomethionine (SeMet) phasing, three mutants (L93M, L146M and L213M) were introduced to facilitate the SeMet labelling, yielding PcoB_Δ27-81_, 3xL/M. The *E. coli* 834(DE3) strain and SelenoMet™ medium (Molecular Dimensions Limited) were used for the SeMet labelling protein expression. The PcoB_Δ27-81_, 3xL/M plasmid was transformed into the *E. coli* 834(DE3) strain (a gift from LP3) and single colonies were inoculated in 5 mL LB medium supplemented with 100 mg/mL ampicillin at 37°C for 8 h. The cells were pelleted and washed 3 times in 1 mL of sterile water, resuspended in 1 mL of sterile water and cultured at 37°C for 16 h in 100 mL minimal media containing L-methionine. Next, the cells were pelleted and washed 3 times in 100 mL sterile water, resuspended in 1 mL water, transferred into 1 L minimal media containing L-SeMet, cultured for 2 h at 30°C, and then induced with 1 mM IPTG at 30°C for 6 h.

The cells were harvested by centrifugation at 8000x*g*, suspended in Tris buffer (20 mM Tris, pH=8, 500 mM NaCl, 10 % v/v Glycerol) and disrupted using sonication. The cell lysate was centrifuged at 165,000x*g* in a Beckman ultracentrifuge (45 Ti rotor, Beckman) for 1 h and the membrane fraction was resuspended in the same Tris buffer containing 1 % w/v N-laurysaccide, and then stirred at 18 °C for 1 h. Subsequently, the outer membrane was pelleted by ultracentrifugation at 165,000x*g* for 1 h. The outer membrane pellets were solubilized in Tris buffer containing 2 % Elugent (Calbiochem) at 4°C for 16 h with stirring. The extract was centrifuged at 190,000x*g* for 30 min and the supernatant was loaded to 5 mL His-trap column (Cytiva) equilibrated with 30 mL Tris buffer containing 0.05 % w/v lauryldimethylamine N-oxide (LDAO). Then, the column was washed with 6 column volume (CV) equilibration buffer containing 50 mM imidazole. The protein was eluted with equilibration buffer containing 300 mM imidazole. The eluted protein fractions were pooled, concentrated and buffer exchanged using Ultra-15 centrifugal concentrators (Amicon) with 50 kDa MW cut-off to remove excess imidazole. Next, TEV protease was added with a molar ratio of 1:10 to remove the His tag through incubation at 4°C for 16 h. Subsequently, the cleaved sample was loaded to a pre-equilibrated 5 mL His-trap column using equilibration buffer. The tag removed protein sample was collected in flow-through and eluted with equilibration buffer containing 40 mM imidazole. Desired fractions were concentrated and buffer exchanged using 50 kDa MW cut-off concentrators (Amicon) to size-exclusion chromatography (SEC) buffer 20 mM Tris, pH=8.0, 100 mM NaCl, 5 % w/w Glycerol and 0.4 % w/w C8E4 detergent. As a polishing step, the protein was finally SEC purified (Superdex 200 10/300; GE Healthcare) and the purified protein was concentrated to 10 mg/mL for the crystallization.

### Crystallization

Initial crystallization for native PcoB and SeMet-PcoB was performed using MemGold, MemGold2, Memstar and MemSys screens from Molecular Dimensions by sitting-drop vapor diffusion using a Mosquito robot at the Lund protein production platform (LP3) by mixing 0.2 μL protein sample with 0.2 μL reservoir solution. The initial hits appeared following two days in a condition with 8 % w/v PEG4000, 0.4 M NaSCN, 0.1 M sodium acetate, pH=4.0, and was optimized using grid-screening and using larger hanging-drop vapor diffusion drops. The best native PcoB crystals grew in 10 % w/v PEG4000, 0.8 M NaSCN, 0.1 M sodium acetate pH=4.0. SeMet-PcoB crystals were obtained from in 30 % w/v PEG400 and 0.1 M sodium citrate, pH=4.0. The crystals were cryoprotected, harvested and flash-cooled in liquid nitrogen for data collection at SLS.

### Data collection and structure determination

Native and SeMet PcoB X-ray diffraction data was collected at the Paul Scherrer Institute, Switzerland on beam line X06SA (SLS). The data was processed and scaled using the software XDS (38). The crystals belonged to space group C2221 with cell parameters a = 64.489 Å, b = 75.538 Å, c = 91.51 Å, α=β=γ=90°. The SeMet data was collected at the wavelength 0.9798 Å. Initial crystallographic phases were calculated by Phenix AutoSol using SAD phasing (39). 7 Selenium atoms were located and refined, then Autobuild (40) was performed to generate the initial model. The initial structure was employed as starting model to determine the PcoB structure of high-resolution native dataset. Model building was performed manually in Coot (41) and refined by phenix.refine (42) in iterative steps. 198 residues were built and refined in the final structure with 95.85 % of residues in the favoured region of the Ramachandran plot (**Table 1**). The final model displayed Ramachandran favored, allowed and outliers of 95.88, 4.12 and 0.00%, respectively. Rotamer outliers were 0.00% and the clash-score was 3.72.

### Cu^2+^ binding stoichiometry determination with ICP-MS

Cu2^+^ binding stoichiometry to wild type PcoB and mutants was measured by inductively coupled plasma mass spectrometry (ICP-MS). The protein forms, purified as mentioned above, were diluted to 3-10 μM with buffer containing 50 mM Tris, pH=8.0, 500 mM NaCl, 10 % w/v glycerol, and 0.05 % w/v LDAO, or 20 mM Tris, pH=8.0, 500 mM NaCl, 10 % w/v glycerol, and 0.8 % w/v OG. 2-5 molar equivalents of CuCl_2_ were subsequently added to the protein solution and incubated at 18 °C for 15 min and excess Cu^2+^ was removed with 5 mL HiTrap desalting columns (Cytiva) packed with Sephadex G-25 resin, pre-equilibrated with the respective buffer. Following elution, the protein concentration was determined by a Bradford assay using BSA as standard. For ICP-MS sample preparation, eluted protein samples were digested in 8 % v/v HNO3 at 80 °C for 12 h. Samples were subsequently diluted to adjust the HNO3 concentration to 3 % and ICP-MS was performed with a Hewlett-Packard 4500 ICP mass spectrometer (Agilent Technologies) connected to a CETACASX-500 auto-sampler for sample injection. Control experiments were conducted in the same buffer excluding the protein for background correction. For the binding experiment high-purity TraceSelect nitric acid, H2O and ICP-MS standards were purchased from Sigma-Aldrich.

### Proteoliposome Cu^2+^ flux assay

PcoB forms were evaluated functionally in liposomes referred to as small unilamellar vesicles (SUVs). Following purification, wildtype PcoB and different mutants were reconstituted in liposomes with a lipid composition of *E. coli* polar lipids (Avanti, US) and 1-palmitoyl-2-oleoyl-sn-glycero-3-phosphocholine (POPC) (Sigma Aldrich) in a 3:1 molar ratio. The lipids were dissolved in chloroform in a glass vial to a concentration of 25 mg/mL followed by treatment using N2 to form a thin lipid bilayer. The lipid film was kept under a N2 stream for 2 h to achieve complete chloroform removal. The lipid film was rehydrated with reconstitution buffer (20 mM Tris, pH=7.4, 200 mM NaCl) with added fluorophore, 10 mM Pyranine (Sigma), to a concentration of 20 mg/mL lipids. The lipid suspension was applied to a sonication bath for 3 times x 15 min, with a 5 min break in-between the cycles. Next, the lipids were frozen in liquid nitrogen and thawed three times, and then the lipids were passed through a 100 nm polycarbonate filter 11 times, using an extruder (Mini-Extruder, Avanti). The lipids were diluted to 4 mg/mL with reconstitution buffer containing 25 % v/v glycerol and 1 % w/v OG, followed by gradual addition of 0.2 % w/v Triton X-100. PcoB and mutants were added to each their lipid suspension using a lipid-to-protein-ratio (LPR) of 20 and each sample was dialyzed for 16 h at 4 oC in reconstitution buffer. The samples were centrifuged at 57,000 xg for 1.5 h and the resulting pellets were suspended in reconstitution buffer. Traditionally, Cu2^+^ flux assays are measured using fluorophores such as Fluozin-1 or −3, which are being activated by addition of Zn^2+^ and quenched by addition of Cu2^+^. However, upon addition of Zn^2+^ and Cu2^+^ the PcoB protein appeared to degrade, in contrast to supplementation of Cu2^+^ only, where only a single band is present on SDS-PAGE analysis. Consequently, we employed fluorescent dye pyranine, which has previously proven effective in measuring the flux of Cu2^+^ ions (43–45). The assay was performed on an SX-20 Stopped-Flow Spectrometer system (Applied Photophysics) equipped with a 495 bandpass filter, where the liposomes were mixed with reaction buffer with 0.1 mM CuCl_2_. Data were collected at 510 nm at a 90° angle for 10 s. All data were collected at 18 oC. Empty liposomes were used as a negative control. The data were analyzed in Pro-Data Viewer (Applied Photosystems) and plotted in GraphPad Prism (v. 9.1.2). Data for each sample were the average of 5 readings. Data were fitted using a double exponential fit. The smallest rate constant is unaffected by changes in PcoB reconstitution efficiency, while the larger rate constant corresponds liposomes containing PcoB. The rates where then adjusted using the wildtype experiments (equivalent to 100 % activity). The reconstitution experiments were performed three times to achieve data reproducibility. Error bars denote the k-rate standard deviation (SD) between the three separate reconstitutions (N = 3) for each construct. SD was calculated using GraphPad Prism.

### MD simulation system design and analyses

The asymmetric lipid bilayer was built using the membrane builder (46) in CHARMM-GUI (47) with the inner leaflet containing 1,1’-palmitoyl-2,2’-vacenoyl cardiolipin (PVCL2), 1-palmitoyl(16:0)-2-palmitoleoyl(16:1 cis-9)-phosphatidylethanolamine (PPPE) and 1-palmitoyl(16:0)-2-vacenoyl(18:1 cis-11)-phosphatidylglycerol (PVPG) lipids while the outer leaflet contained lipopolysaccharides (LPS) with lipid A of type 1 tail and core R1 and a 1 o-antigen polysaccharide chain (48). The protein was inserted into the membrane based on prediction from the Orientations of Proteins in Membranes (OPM) (49). Predicted pKa values were calculated with the Propka-3.1 program (50) and residues Glu130, Glu187, Glu207 and Glu243 were protonated. The system was simulated using the GROMACS 2019 MD simulation engine (51) with the CHARMM36 all-atom force field (52, 53). The system contained 25,465 TIP3P water molecules, 550 Na-ions and 47 Cl-ions (in total 131,956 atoms). A 5,000-step energy minimization was followed by a 30 ns equilibration run during which the protein backbone, side chain, lipid, and water atoms were successively unrestrained in six consecutive 5-ns steps leading to the 500 ns production run. The simulation time step was 2 fs and a Parrinello-Rahman semi-isotropic pressure coupling (54, 55) with a compressibility of 4.5e-5 bar^-1^ was applied with a coupling constant of 1.0 at 1 bar and the temperature was maintained at 303.15 K using the Nose-Hoover temperature coupling (56, 57). A Cu^2+^ ion was added to the final frame of the production run of 500 ns by replacing a Na^+^ ions at the outer membrane entrance and reparametrizing according to Cu^2+^ parameters from (58). The Cu^2+^ ion was pulled from the outer to inner leaflet using the Gromacs pull code. The Cu^2+^ ion was pulled for 3.5 ns at a pulling rate of 1.5 nm/ns at a harmonic force of 1000 kJ/mol/nm^2^. The Na and Cl-ions were restrained during pulling to avoid interference with ion-entry dynamics.

## Acknowledgments

We are grateful for the staff at the Lund Protein Production Platform (LP3) for providing the *E. coli* 834(DE3) strain and for assistance with the early crystallization experiments. We also appreciate the help with the crystal screening and data collection at the Swiss Light Source (SLS), the Paul Scherrer Institute, Villigen, beam line X06SAWe acknowledge Zhila Nikrozi and Klaus Qvortrup from the Core Facility for Integrated Microscopy (CFIM), Faculty of Health and Medical Sciences, University of Copenhagen for the help of preparing fixed bacteria TEM sample preparation and imaging, and Prof. Michael Davies for access to stop-flow cytometry equipment. Access to synchrotron sources was supported by The Danscatt program of The Independent Research Fund Denmark. GM is supported by the National Institute of General Medical Sciences of the National Institutes of Health (R35GM128704) and by the Robert A. Welch foundation (AT-1935-20170325; AT-2073-20210327). PG is supported by the following Foundations: Lundbeck, Knut and Alice Wallenberg, Carlsberg, Novo-Nordisk, Brødrene Hartmann, Agnes og Poul Friis, Augustinus, Crafoord as well as The Per-Eric and Ulla Schyberg. Funding is also obtained from The Independent Research Fund Denmark, the Swedish Research Council and through a Michaelsen scholarship. DRM was funded by Carl Tryggers foundation (CTS 17:22), MA was funded by a Swedish Research Council Starting Grant (2020-03840). The computations were performed on resources provided by the Swedish National Infrastructure for Computing (SNIC) through the High-Performance Computing Center North (HPC2N) under project SNIC 2018/2-32 and SNIC 2019/2-29. The funders had no role in study design, data collection and analysis, decision to publish, or preparation of the manuscript. The content is solely the responsibility of the authors and does not necessarily represent the official views of the National Institutes of Health.

## Author Contributions

P.L. and P.G. identified and developed the project

P.L. crystallized PcoB

P.L. and P.G. solved the structure

P.L. refined the structure

P.L., N.N., K.G., J.W.M., K. L-P and P.G. analyzed the structure

N.N. purified constructs for the ICP-MS and liposome experiments

S.A. and G.M. performed ICP-MS experiments

N.N., E.R.B., J.W.M. performed liposome experiments

J.W.M. performed the EVcouplings analysis

D.R.M. and M.A. performed and analyzed MD simulations

P.L., N.N. and K.G. wrote the initial draft

P.L., N.N., K.G., J.W.M. and P.G. finalized the manuscript with comments from all authors

## Data deposition

The structure of PcoB_Δ32-82_ reported in this paper will be released by the Protein Data Bank upon acceptance of this manuscript (PDB, PDB-ID 7PGE).

---

**Figure 3 - figure supplement 1.**
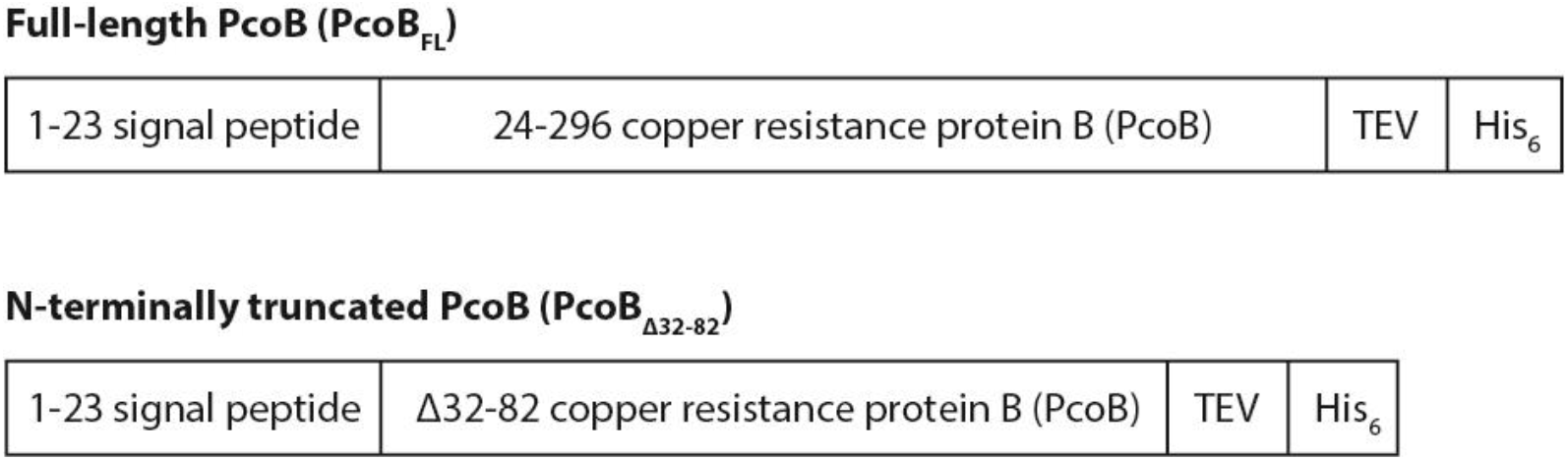
Construct design of PcoB.

**Figure 3 - figure supplement 2.**
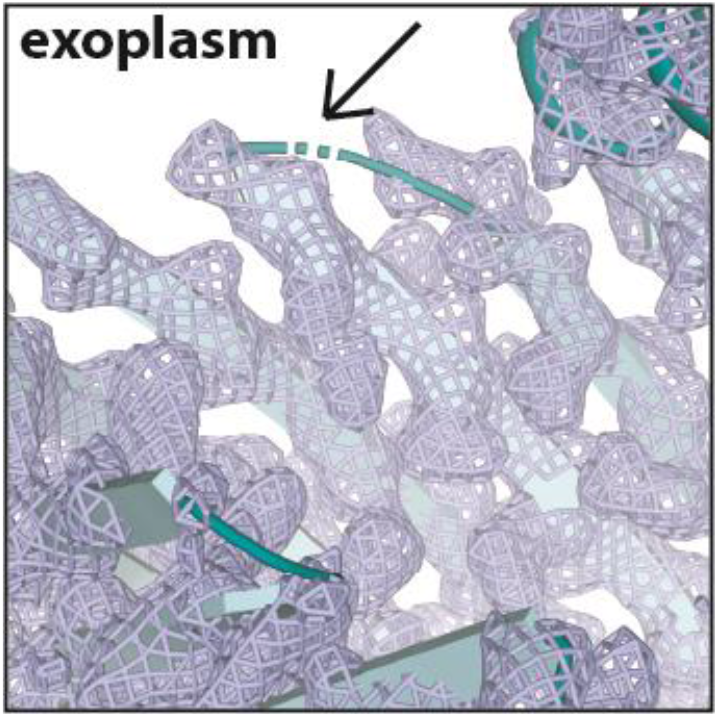
Details of the structure of PcoB. The disordered loop region in the PcoB structure (cyan). The final 2Fo-Fc electron density is shown in blue mesh, σ = 1.0. Arrow points to the unmodeled loop (dashed line).

**Figure 3 - figure supplement 3.**
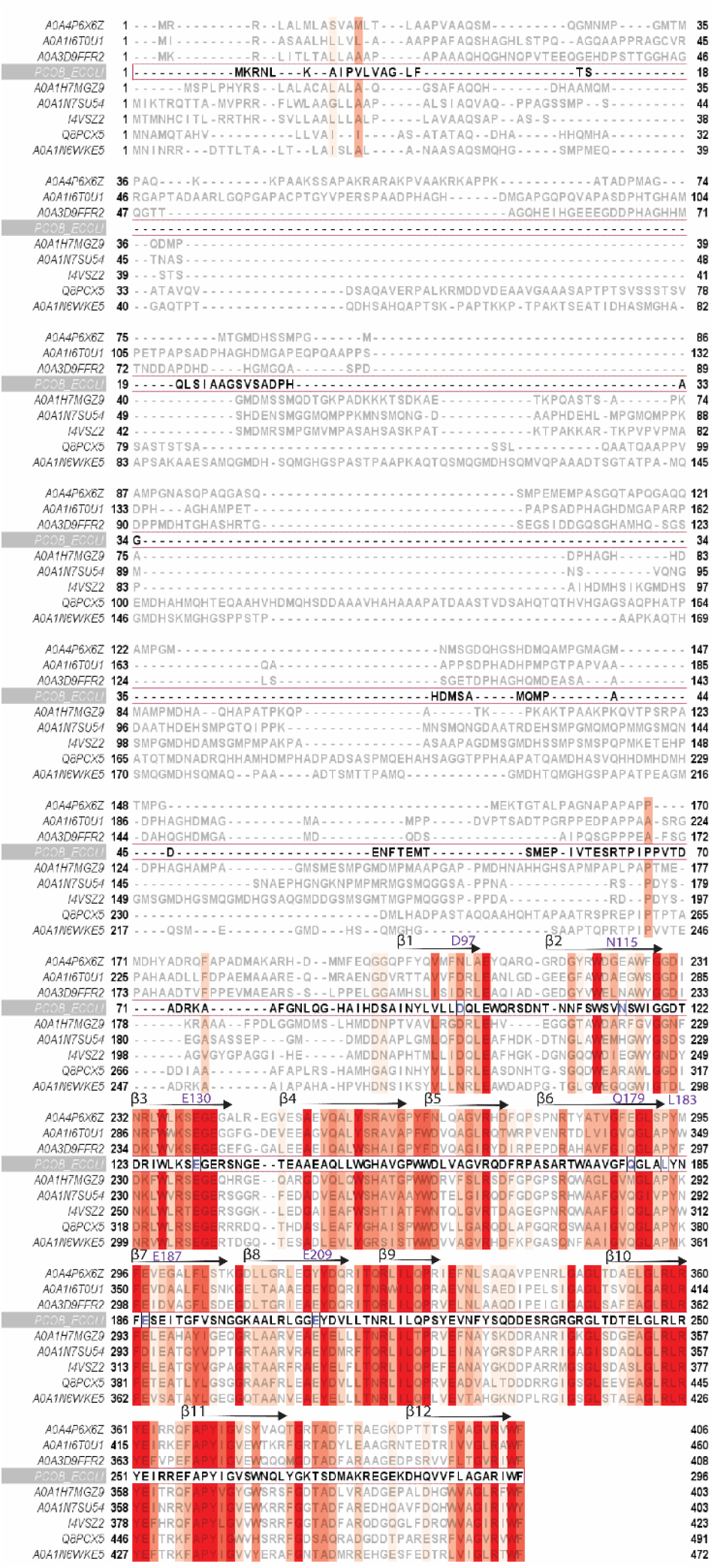
Sequence conservation among PcoB proteins. Accession numbers refer to Uniprot with *E. coli* PcoB highlighted in bold. Red columns indicate the most conserved residues. Structurally and functionally important residues are shown in purple with the residue number of *E. coli* PcoB indicated above each row. The location of the β-strands of the structure is shown. The alignment was generated through a Uniprot Blastp search using the *E. coli* PcoB as a template, thereby securing 250 proteins. These sequences were aligned using Clustal Omega and sequence redundancy (higher than 75 % identity) was removed using Jalview.

**Figure 4 - figure supplement 1.**
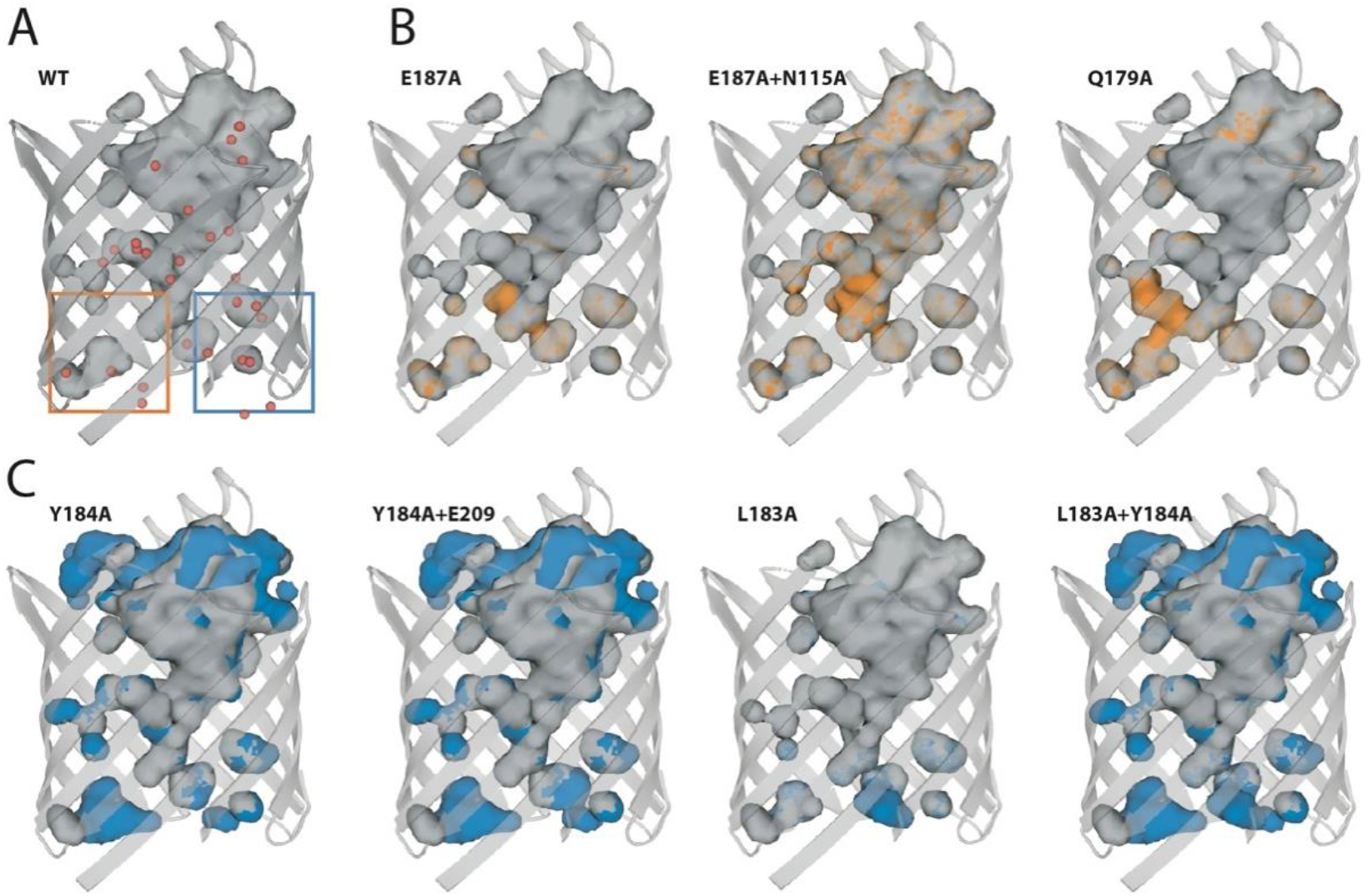
Two putative exit paths in PcoB. **(A)**The structurally determined wild-type PcoB (gray) does not provide a continuous pore as shown using the surface of internal cavities (grey) and crystal waters (red spheres). *In silico* analysis was conducted to assess if mutant forms may render the pore more open. **(B)**Mutations related to the proposed exit pathway (orange). From left to right: Glu187Ala; Glu187Ala and Asn115Ala; Gln179Ala. **(C)**From left to right: Tyr184Ala; Tyr184Ala and Glu209Ala; Leu183Ala; Leu183Ala and Tyr184Ala.

**Figure 6 - figure supplement 1.**
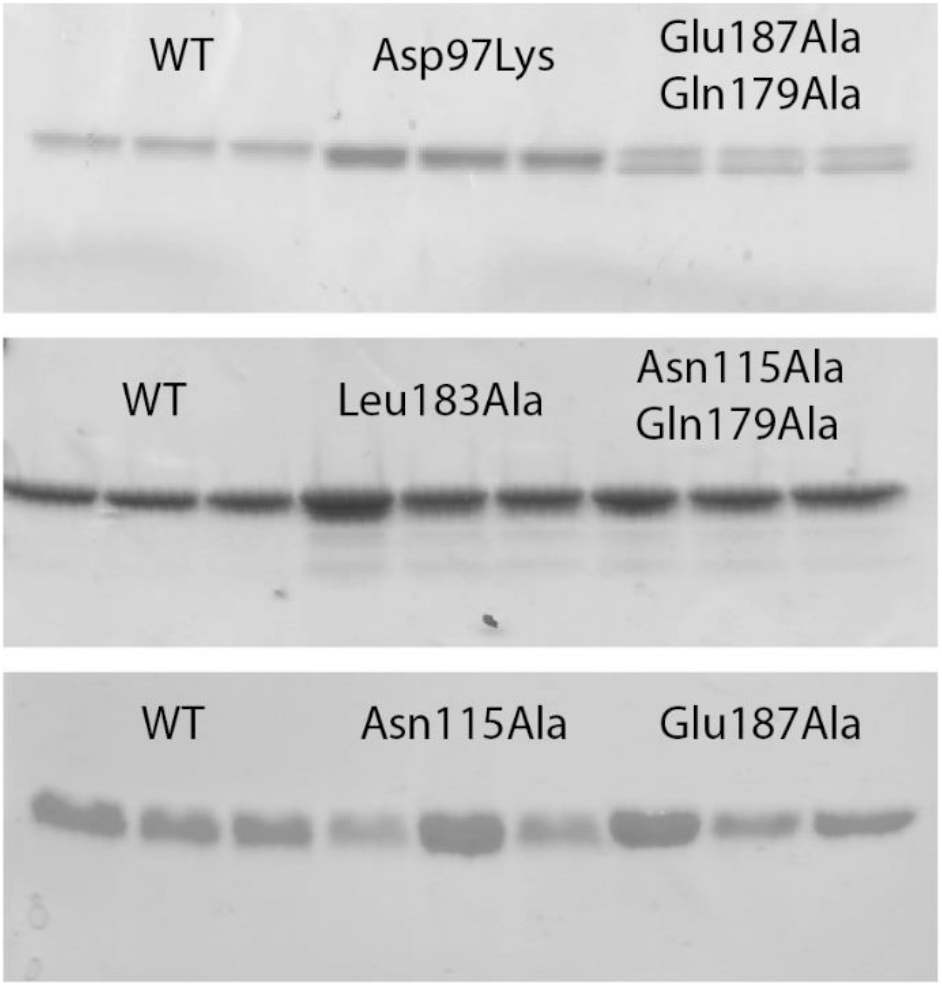
SDS-PAGE gels for various PcoB mutants. SDS-PAGE used for analyses of all PcoB forms reconstituted into liposomes (**Figure 6 - table supplement 1**). All mutations were quantified against wild-type (WT). The double mutants and Leu183Ala showed degrees of degradation, and were excluded from the final analysis.

**Figure 6 - figure supplement 2.**
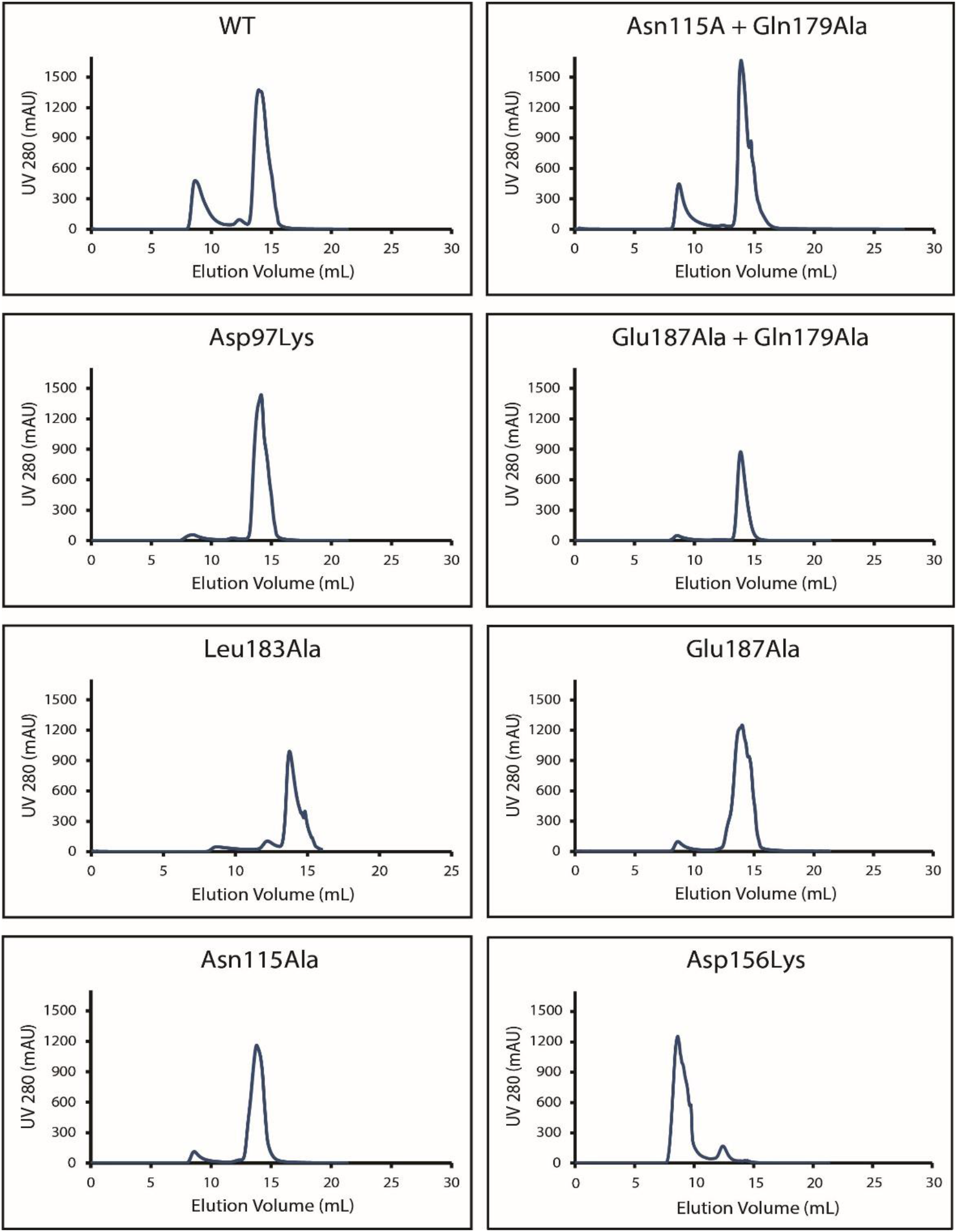
Size-exclusion chromatography profiles of the studied PcoB forms. His-tagged cleaved samples were injected into an pre-equilibrated Superdex 200 Increase 10/300 GL column mounted on an Äkta Avant system. The employed buffer included 20 mM Tris-HCl pH=8, 100 mM NaCl, 5 % Glycerol, 0.8% OG and the flow rate was 0.4 mL/min for all forms.

**Figure 6 - table supplement 1.** Quantification constants of the three separate reconstitutions using ImageJ of the SDS-PAGE. SDS-PAGE quantification analyses of all PcoB forms reconstituted in liposomes. All mutations were quantified against the first wild-type (WT) lane (**Figure 6**, **Figure 6 - figure supplement 1**, **Figure 6 - source data 1**) normalized to 1.0. The analysis was performed by plotting the band intensities in ImageJ and taking the integral from these separate plots. The factors were then divided by the integral from the WT lane 1, giving the constants in this table, of which the relative activity could be adjusted (Figure 6B).

|  | <u>WT</u> | <u>ASP97LYS</u> | <u>ASN115ALA</u> | <u>GLU187ALA</u> |
| --- | --- | --- | --- | --- |
| <u>CONSTANT #</u> |  |  |  |  |
| 1 | 1.0 | 2.2 | 0.7 | 1.0 |
| 2 | 1.1 | 2.2 | 1.6 | 0.8 |
| 3 | 1.1 | 2.2 | 0.9 | 1.5 |

